# Efficient base-catalysed Kemp elimination in an engineered ancestral enzyme

**DOI:** 10.1101/2022.07.28.501888

**Authors:** Luis I. Gutierrez-Rus, Miguel Alcalde, Valeria A Risso, Jose M. Sanchez-Ruiz

## Abstract

The routine generation of enzymes with completely new active sites is one of the major unsolved problems in protein engineering. Advances in this field have been so far modest, perhaps due, at least in part, to the widespread use of modern natural proteins as scaffolds for *de novo* engineering. Most modern proteins are highly evolved and specialized, and, consequently, difficult to repurpose for completely new functionalities. Conceivably, resurrected ancestral proteins with the biophysical properties that promote evolvability, such as high stability and conformational diversity, could provide better scaffolds for *de novo* enzyme generation. Kemp elimination, a non-natural reaction that provides a simple model of proton abstraction from carbon, has been extensively used as a benchmark in *de novo* enzyme engineering. Here, we present an engineered ancestral β-lactamase with a new active site capable of efficiently catalysing the Kemp elimination. Our Kemp eliminase is the outcome of a minimalist design based on a single function-generating mutation followed by sharply-focused, low-throughput library screening. Yet, its catalytic parameters (k_cat_/K_M_=2·10^5^ M^−1^s^−1^, k_cat_=635 s^−1^) compare favourably with the average modern natural enzyme and with the best proton-abstraction *de novo* Kemp eliminases reported in the literature. General implications of our results for *de novo* enzyme engineering are discussed.

## 1. Introduction

In the early 1960s, Linus Pauling and Emile Zurkenkandl published two papers that played a crucial role in the emergence and subsequent development of the molecular evolution field. In the first paper [1], they introduced the molecular clock hypothesis and, hence, the possibility of using sequences of homolog proteins to estimate species divergence times. In the second paper [2], they proposed that plausible approximations to the sequences of ancestral proteins can be derived from suitable analyses of the known sequences of their modern counterparts. Ancestral sequence reconstruction has been amply used in the post-genomic era as a tool to address fundamental problems in molecular evolution[3-5]. Furthermore, it has been found that ancestral proteins “resurrected” in the lab (*i.e*., the proteins encoded by the reconstructed sequences) often display interesting and even extreme properties [6-16] plausibly reflecting ancestral adaptations to unusual intra- and extra-cellular environments. Resurrected ancestral proteins often display high stability, supporting the frequently hypothesized thermophilic nature of ancient life. Also, they often show efficient heterologous expression in modern hosts, possibly reflecting their emergence prior to the availability of efficient cellular folding assistance and, therefore, that, unlike many modern proteins, they do not rely on having adapted to (*i.e*., having co-evolved with) the folding assistance machinery of the host to fold efficiently. Finally, in a significant number of ancestral reconstruction studies, the resurrected enzymes were found to be able to catalyse several related reactions. Such promiscuity, to use the accepted term in the field, may reflect that primordial enzymes were generalists, as proposed by Jensen many years ago [17], or perhaps that the promiscuous resurrected ancestral enzymes corresponded to pre-duplication nodes in the evolution of new functionalities [10].

Beyond specific evolutionary narratives, it is clear that the unusual properties of resurrected proteins may be valuable in biotechnological application scenarios [5,7,12-14,18-22]. Promiscuity, for instance, is most likely linked to conformational diversity, *i.e*., to the fact that the protein exists in solution as an ensemble of more or less related conformations, with different functionalities being linked to different subsets of conformations [23]. It is well known that conformational diversity promotes evolvability, *i.e*., the capability to evolve new functions [24-27]. The reason for this is that, if a few rare conformations are competent for the targeted activity, their population in the ensemble can be enhanced through rational design or, more likely, through standard directed evolution. Enhanced stability, besides being a biotechnologically useful property by itself, it is also known to contribute to evolvability [28], as functionally useful but destabilizing mutations will be accepted in a high stability protein scaffold, while they will likely compromise proper folding in a moderately stable protein.

Furthermore, and contrary to naïve expectations, enhanced conformational diversity and high stability are not necessarily mutually exclusive features [29]. In an ancestral reconstruction exercise targeting the antibiotic-resistance enzyme β-lactamase published about ten years ago [7], we found the resurrected proteins corresponding to ancient Precambrian phylogenetic nodes to be not only highly stable, with denaturation temperatures around 30 degrees above those of their modern mesophilic counterparts, but also promiscuous, being able to degrade a variety of lactam antibiotics, including third-generation. For comparison, TEM-1 β-lactamase, a typical modern β-lactamase, is a penicillin specialist and displays very low levels of activity with other lactam antibiotics. Subsequent experimental and computational studies [30,31] confirmed that the substrate promiscuity of the ancestral β-lactamases was indeed linked to enhanced conformational diversity. Resurrected Precambrian β-lactamases are therefore highly-stable and conformationally diverse to some substantial extent.

Overall, it emerges that resurrected ancestral proteins in general, and our highly stable and conformationally diverse Precambrian β-lactamases in particular, could perhaps provide superior starting points for protein engineering. In order to systematically explore this possibility, we recently started a research program aimed at using our resurrected ancestral β-lactamases as scaffolds for *de novo* enzyme generation, a fundamental unsolved problem in protein engineering [32]. We targeted Kemp elimination (Figure 1), a simple model of a fundamental chemical process, proton abstraction from carbon, and an extensively used benchmark in *de novo* enzyme engineering. Specifically, we aimed at generating efficient Kemp eliminases with minimal design and screening efforts. In a first step [33], we succeeded in using a single-mutation, minimalist design to generate significant levels of Kemp elimination activity in ancestral β-lactamase scaffolds. In the second step [34], we improved these starting activities on the basis of ultra-low-throughput screening computationally focused on the new active site region. Here, we report the third step along these lines. The interactions at the original *de novo* active site have probably been optimized to a substantial extent by our previous efforts. Therefore, we have devised a different approach based on the extension of the protein through the introduction of an additional polypeptide segment near the new active site. In this way, new interactions are generated and can be exploited for *de novo* activity enhancement. In fact, by screening a small library focused at the additional segment, we reach in this work a catalytic efficiency of k_cat_/K_M_=2·10^5^ M^−1^s^−1^, and a turnover number of k_cat_=635 s^−1^. These Michaelis-Menten catalytic parameters compare favourably with those for the average modern natural enzyme [35] and the best proton-abstraction *de novo* Kemp eliminases reported in the literature [36]. General implications of these results for *de novo* enzyme engineering are elaborated in the Discussion section.

**Figure 1.**
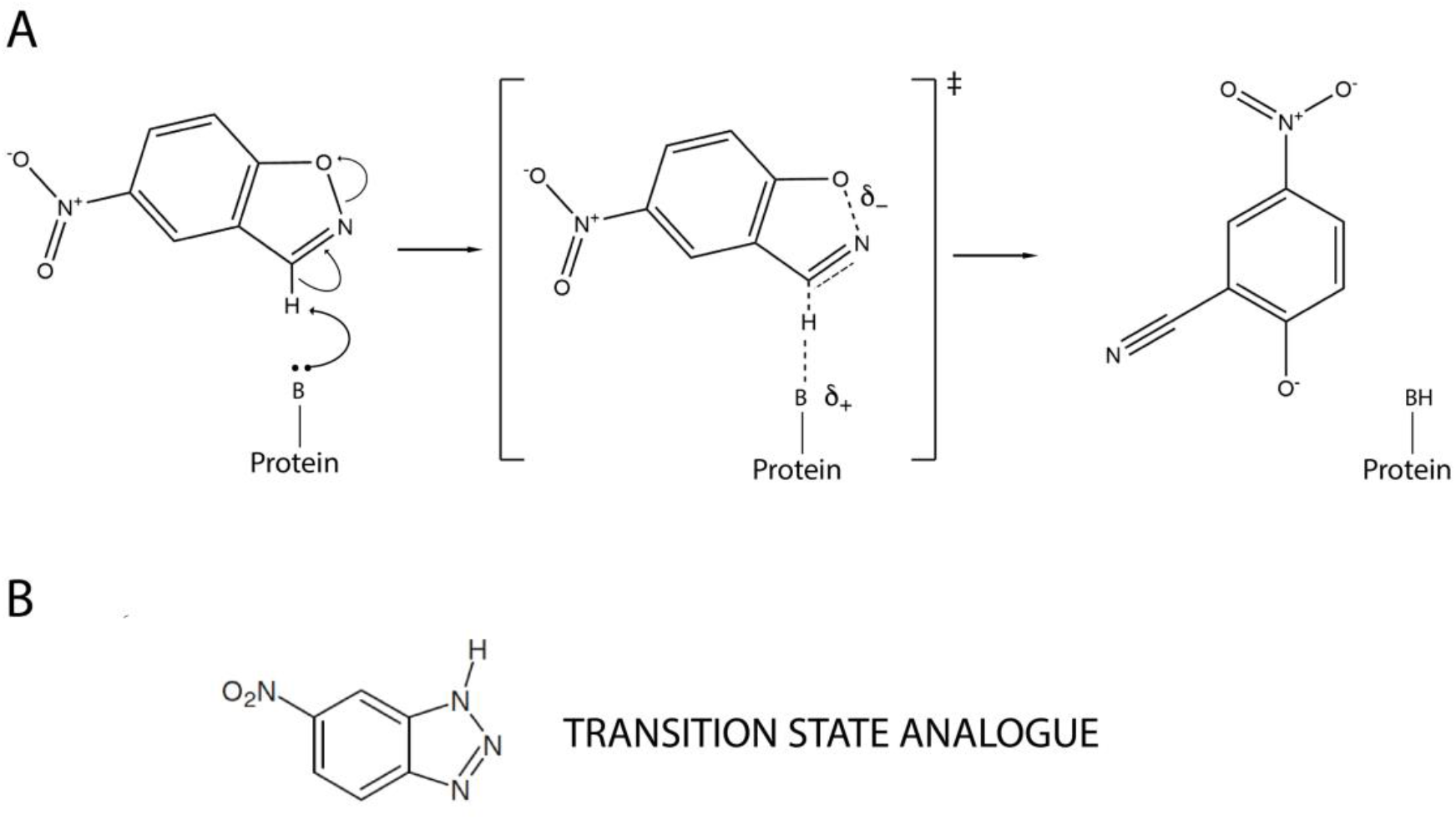
Mechanism of base-catalysed Kemp elimination showing a proposed transition state structure (**A)**. A transition state analogue, 5(6)-nitrobenzotriazole is also shown (**B**).

## 2. Results

### 2.1 Kemp Eliminase Variants Used as Starting Point for This Work

Recently [33], we generated a completely new active site for Kemp elimination, *i.e*., distinct from the antibiotic degradation active site, using a minimalist approach based on a single mutation. Specifically, a hydrophobic-to-ionizable mutation generated both a cavity for substrate binding and a catalytic base capable of performing proton abstraction. Both, the high stability and the conformational diversity of the ancestral β-lactamase scaffolds used likely played a role in the success of the minimalist design [33,37]. The function-generating mutation, a tryptophan to aspartate amino acid replacement at position 229, is clearly disruptive and may lead to folding problems if implemented in a β-lactamase of moderate stability. Furthermore, the cavity produced by the mutation cannot exactly match the shape of the Kemp reactant and substrate binding must rely therefore on local flexibility in the region of the new active site. Indeed, while the minimalist design was successful in a number of resurrected Precambrian β-lactamases, it failed to lead to significant *de novo* activity levels when implemented in ten different modern β-lactamases.

The activity generated by the W229D was found to be enhanced by a second F290W mutation [33], resulting in a catalytic efficiency of about 10^4^ M^−1^s^−1^ with a turnover number of about 10 s^−1^ for Kemp elimination catalysis. Figure 2A shows the 3D-struture of the more active *de novo* Kemp eliminase we initially obtained [33] through a scan of the function-generating W229D mutation on several resurrected ancestral β-lactamases. The structure shown includes a bound transition-state analogue (Figure 1B) which indicates the location of the engineered active site.

**Figure 2.**
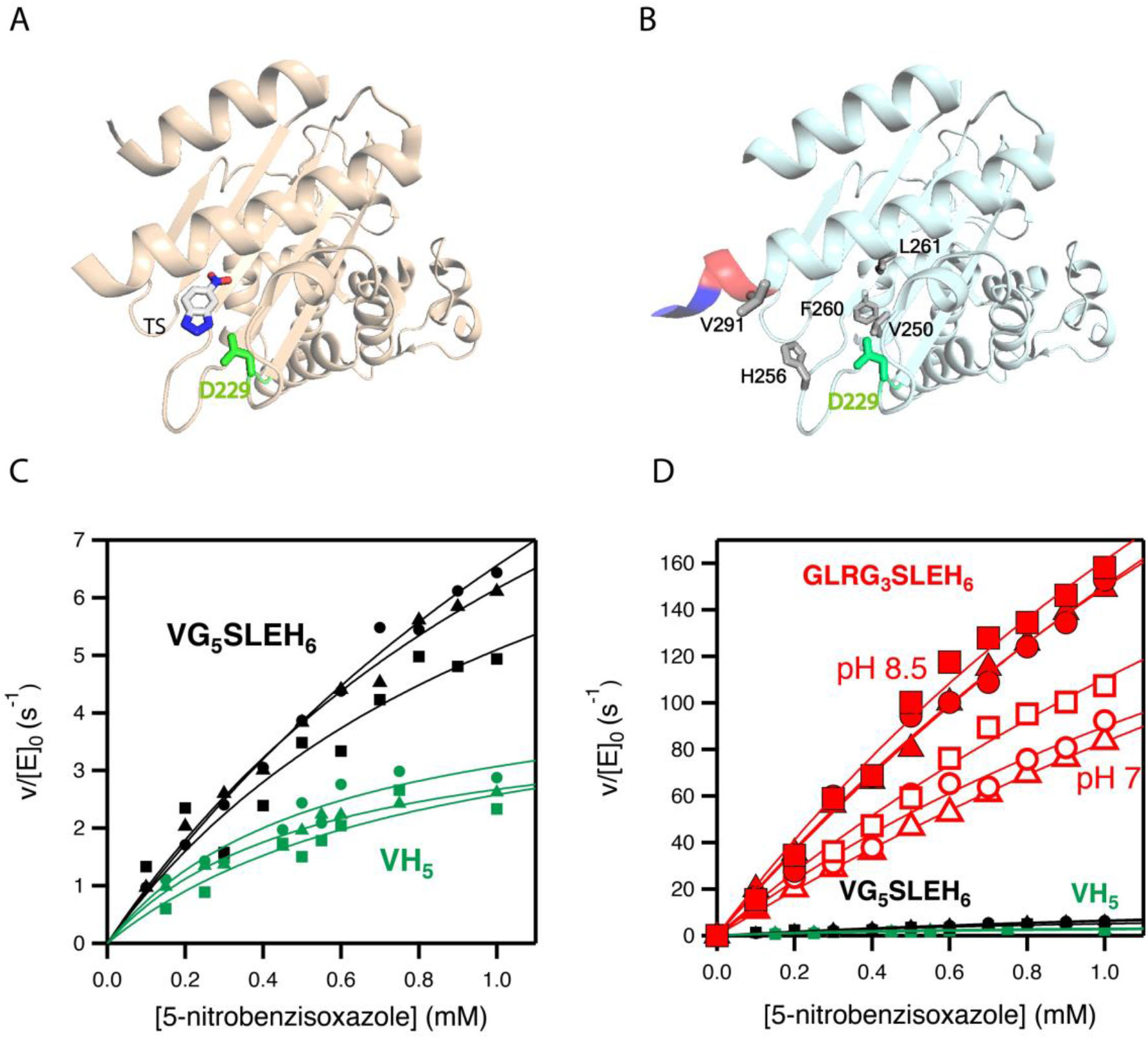
3D-structures and catalytic properties of the Kemp eliminase variants used as starting points for the enzyme engineering reported in this work. (**A**) Structure (PDB ID 5FQK) of the W229D/F290W variant of an ancestral β-lactamase with Kemp eliminase activity [33]. The W229D mutation generates the new function and the catalytic base (D229) introduced by the function-generation mutation is shown. The structure also includes a transition state analogue (see Figure 1), which indicates the location of the new active site. (**B**) The Kemp elimination activity of the protein shown in A could be enhanced by several amino acid replacements at the *de novo* active site region resulting from computationally-focused ultra-low-throughput screening [34]. The activity-enhancing residues are highlighted in grey in the structure shown here (PDB ID 6TXD). The His-tag attached to the carboxyl terminus for purification purposes is also highlighted (red and blue colours). Note that one of the activity enhancing mutations replaces de first histidine with a valine. Therefore, the protein shown has a VH_5_ tail attached to the carboxyl terminus. The background variant used in this work has VG_5_SLEH_6_ attached to the carboxyl terminus, introducing a polypeptide segment between the caxboxyl terminus and the His-tag, but keeping the valine. (**C**) Michaelis-Menten profiles for the proteins with VH_5_ and VG_5_SLEH_6_ attached to the carboxyl terminus. (**D**) The differences in catalysis observed in C are very small compared with the activity enhancement achieved in the screening efforts reported this work (variant with GLRG_3_SLEH_6_ attached to the carboxyl terminus). Note that three Michaelis-Menten profiles have been independently determined for each of the variants shown in C and D.

Subsequently [34] we used computationally-focused screening to targeted at the active site region to further increase the Kemp elimination activity of our best W229D/F290W variant of an ancestral β-lactamase scaffold. The enhancement in catalytic efficiency obtained was moderate, but the turnover number was raised by about one order of magnitude. The mutations introduced at this stage are highlighted in the structure shown Figure 2B. Note that in all our Kemp eliminases the *de novo* active site is located near the carboxyl terminus and that His-tag is routinely attached to the carboxyl terminus residue to enable protein purification by affinity chromatography. Note also that one of the activity-enhancing mutations shown in Figure 2B actually replaces the first histidine residue of the purification tag with a valine.

### 2.2 Combinatorial Library Design and Screening

In this work, we have inserted a polypeptide segment between the carboxyl terminus residue and the His tag (Figure 3A) of our previous best Kemp eliminase. This insertion retains the valine residue at the first position after the original carboxyl terminal residue, while the rest of the inserted segment includes several glycine residues and a serine, following the known sequence design principles for soluble and flexible protein linkers [38]. Overall, the β-lactamase variant used here as starting point for directed evolution is identical to the best Kemp eliminase from our previous study [34], except for the presence of a (Gly)_5_-Ser-Leu-Asp-(His)_6_ segment between the extra valine at the carboxyl terminus and the purification His-tag. This insertion has a small effect on catalysis, as shown by the Michaelis-Menten plots and catalytic parameters shown in Figure 2C. In fact, as anticipated in Figure 2D, the effect of the insertion on catalysis is almost negligible compared with the total activity enhancement achieved in this work.

**Figure 3.**
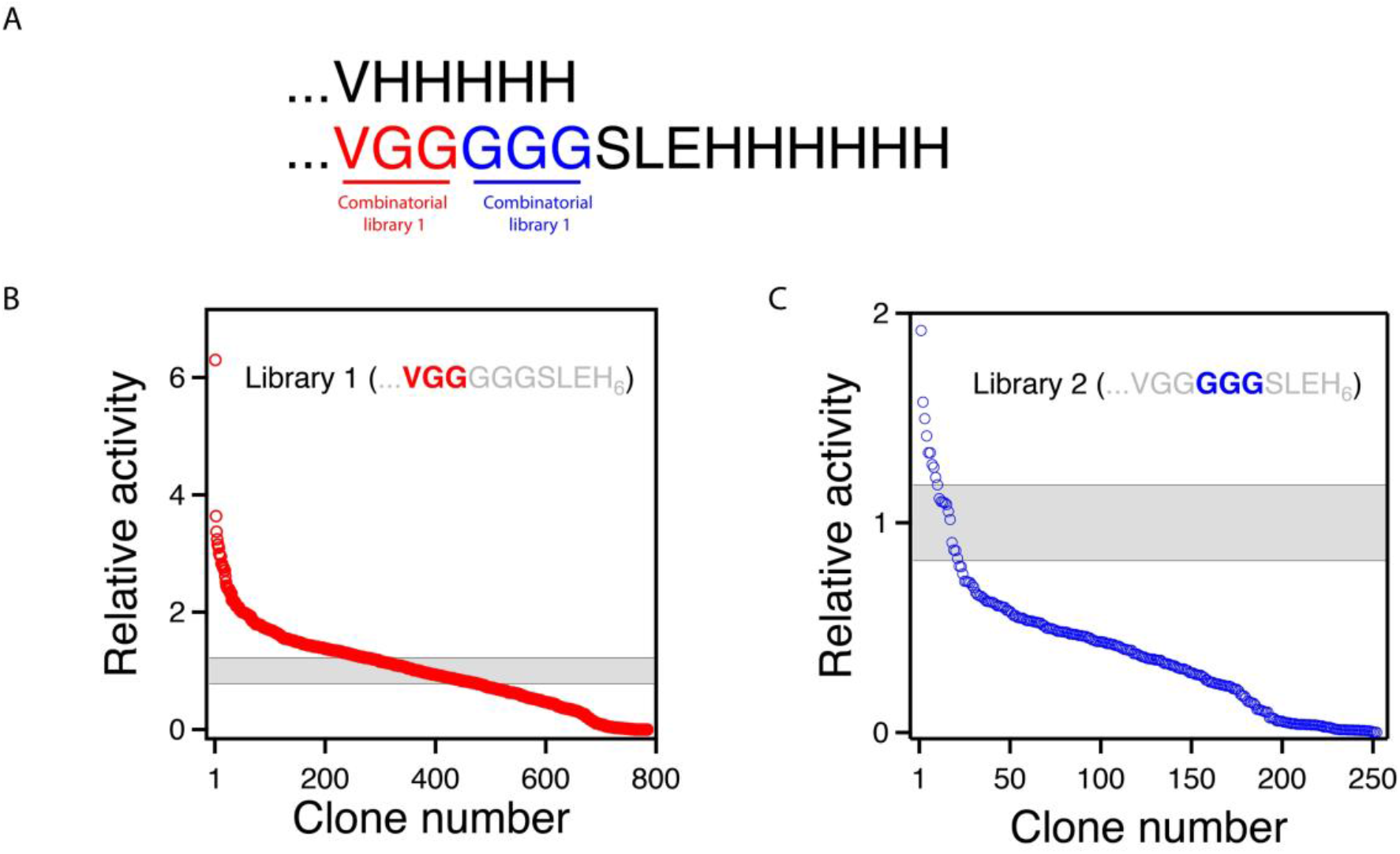
Combinatorial library screening for enhanced Kemp eliminase activity. (**A)** The background variant used for library construction has a VG_5_SLEH_6_ polypeptide attached to the carboxyl terminus. Two 8000-variant combinatorial libraries were prepared including all possible combinations of the 20 amino acids at two sets of the positions, as shown. (**C**) and (**D**) Result of the screening of the two libraries. Clones are ranked according to Kemp eliminase activity. The grey strip represents the average activity of the background variant plus/minus the associated standard error.

The β-lactamase variant described above was used as background for a library comprising all combinations of all possible amino acid residues at the three first positions after the carboxyl terminus (Figure 2C). The rationale behind this approach is that, since the carboxyl terminus is close to the *de novo* active site, some of the library variants may generate interactions that promote catalysis. The combinatorial library spans 20^3^=8000 different amino acid sequences, and about 800 variants were screened for Kemp elimination activity as described in Methods. In this primary screening, most variants displayed significant levels of Kemp elimination activity (Figure 3B), supporting that few of the mutations were disruptive and prevented proper folding. This is of course consistent with the fact that, in this case, library screening samples the sequence space associated with a conformationally flexible polypeptide segment that it is expected to remain largely exposed to the solvent in most cases. Still, a few of the variants were found to display substantially enhanced levels of catalysis for the Kemp elimination reaction (Figure 3B). Since catalysis relies on decreasing the activation free energy of the reaction, it appears reasonable to assume that these variants can generate interactions that stabilize the transition state (*i.e*., the chemical species at the top of the free energy barrier) for Kemp elimination at the *de novo* active site.

### 2.3 Stability and Catalytic Parameters for the Improved Kemp Eliminases

The 7 top variants of the primary screening described above were purified and their Michaelis-Menten profiles of rate *versus* substrate concentration were determined at pH 7 (Figure 4A and Supplementary Figure S1). This secondary screening confirmed the results of the primary screening, as all the selected variants showed substantially enhanced catalysis with respect to the library background (Figure 4A and Table 1). In particular, the variant with the sequence GLR at the three targeted positions shows an enhancement in catalytic parameters of about one order of magnitude over the library background. Note that most of the activity determinations reported in this work have been performed at pH 7. However, it is known that the activity of Kemp eliminase enzymes based on the proton-abstraction mechanism may increase at basic pH, reflecting the deprotonation of the amino acid residue that gives rises to the catalytic base (the aspartate at position 229 in this case). Accordingly, we have determined the profile of rate *versus* substrate concentration for the best GLR variant at pH 8.5. A significant increase in activity with respect to pH 7 is observed (Figure 2D) leading to a catalytic efficiency of k_cat_/K_M_∼2·10^5^ M^−1^s^−1^ and a turnover number of k_cat_∼700 s^−1^ (Table 1).

**Figure 4.**
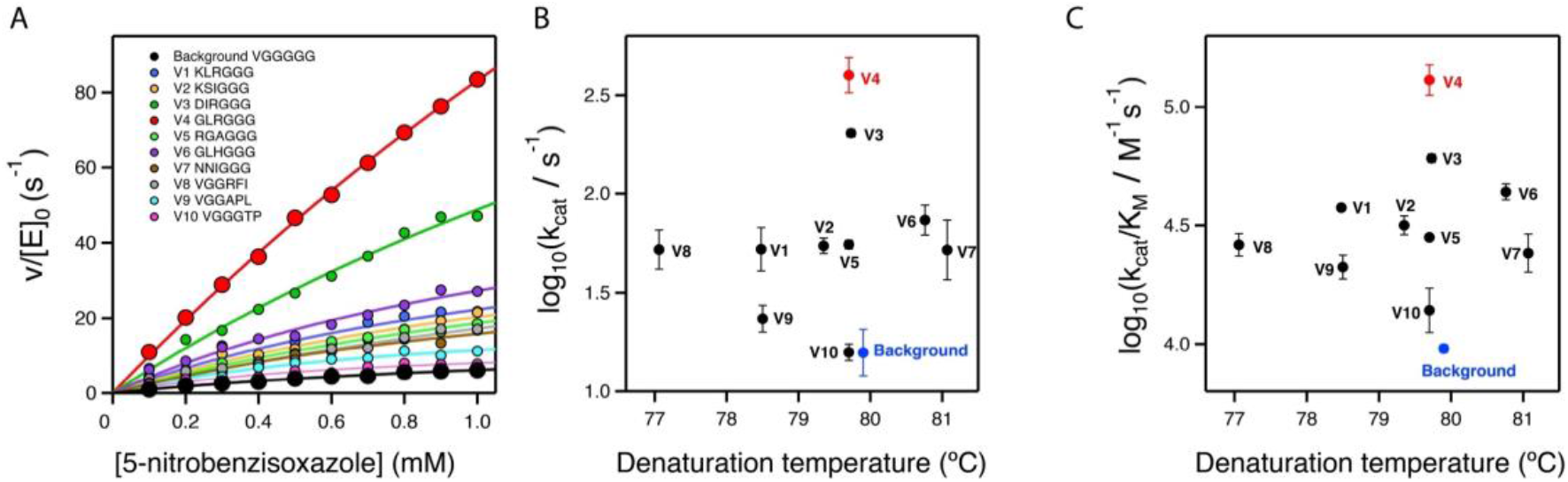
Secondary screening of the top variants from the primary library screening shown in figure 3. (**A)** Michaelis-Menten profiles for the top variants at pH 7 (profiles for the best variant at pH 8.5 are shown in Figure 2B). The sequences at the relevant section of the included polypeptide are shown. Note that, for all variants, three independent Michaelis-Menten profiles were determined (see Supplementary Table S1 and Supplementary Figure S1), although, for the sake of clarity, only one representative profile for each variant is shown here. (**C**) and (**D**) Plots of Michaelis-Menten catalytic parameters for all the variants *versus* denaturation temperature, as determined by differential scanning calorimetry. These plots are scattergrams, indicating the absence of a significant stability/activity trade-off.

**Table 1.** Catalytic parameters for the cleavage of 5-nitrobenzisoxazole at pH 7 (HEPES 10 mM NaCl 100 mM) and 1% acetonitrile and 25 ºC catalyzed by the engineered and evolved versions of Precambrian β-lactamases. For the best variant, data at pH 8.5 are also included.

| Variant | Sequence<sup>a</sup> | $k_{\text{cat}}$ (s<sup>-1</sup>)<sup>b</sup> | $K_M$ (mM)<sup>b</sup> | $k_{\text{cat}}/K_M$ (s<sup>-1</sup> M<sup>-1</sup>)<sup>b</sup> | $T_m$ (C°)<sup>c</sup> |
| --- | --- | --- | --- | --- | --- |
| V4 at pH 8.5 | ...GLRGGG... | 635.0 ± 59.1 | 3.14 ± 0.49 | (2.0 ± 0.1) $\times 10^5$ | -- |
| V4 | ...GLRGGG... | 407.5 ± 76.7 | 3.35 ± 0.91 | (1.3 ± 0.2) $\times 10^5$ | 79.7 |
| V3 | ...DIRGGG... | 202.9 ± 8.9 | 3.35 ± 0.21 | (6.1 ± 0.3) $\times 10^4$ | 79.7 |
| V6 | ...GLHGGG... | 74.7 ± 13.5 | 1.74 ± 0.46 | (4.4 ± 0.3) $\times 10^4$ | 80.8 |
| V1 | ...KLRGGG... | 54.1 ± 13.2 | 1.45 ± 0.37 | (3.6 ± 0.1) $\times 10^4$ | 78.5 |
| V2 | ...KSIGGG... | 54.7 ± 4.8 | 1.75 ± 0.32 | (3.2 ± 0.3) $\times 10^4$ | 79.4 |
| V5 | ...RGAGGG... | 55.4 ± 2.7 | 1.97 ± 0.1 | (2.82 ± 0.01) $\times 10^4$ | 79.7 |
| V8 | ...VGGRFI... | 53.5 ± 11.0 | 2.10 ± 0.61 | (2.6 ± 0.3) $\times 10^4$ | 77.0 |
| V7 | ...NNIGGG... | 55.4 ± 20.8 | 2.50 ± 1.43 | (2.5 ± 0.4) $\times 10^4$ | 81.1 |
| V9 | ...VGGAPL... | 23.7 ± 3.8 | 1.12 ± 0.16 | (2.1 ± 0.2) $\times 10^4$ | 78.5 |
| V10 | ...VGGGTP... | 15.9 ± 1.5 | 1.19 ± 0.32 | (1.4 ± 0.3) $\times 10^4$ | 79.7 |
| BACKGROUND | ...VGGGGG... | 16.3 ± 4.4 | 1.71 ± 0.48 | (9.6 ± 0.4) $\times 10^3$ | 79.9 |

<sup>a</sup> The sequences of each variant at the randomly mutagenized regions in the libraries are shown. Variants are ranked according to the turnover number. All values given correspond to pH 7, with the exception of the best variant (V4) for which data at pH 7 and pH 8.5 are given. <sup>b</sup> The values shown for the Michaelis-Menten catalytic parameters are the average values from three independent replicates (n = 3 independent determinations of the Michaelis-Menten profiles). The associated standard errors are also given. See Supplementary Figure S1 Supplementary Table S2 for the results of the analysis of the individual profiles. <sup>c</sup> Denaturation temperatures determined by differential scanning calorimetry (DSC) are shown. The error associated with $T_m$ determination by DSC is typically smaller than one degree.

It is interesting that the achieved improvements in catalysis does not come with a significant cost in stability, as shown by the denaturation temperature values determined by differential scanning calorimetry (Figures 4B and 4C, and Table 1). The lack of a substantial activity-stability trade-off can be put down to the fact that our engineering approach does not manipulate already-existing interactions, but introduces new ones.

### 2.4 Structural Analysis of the Catalysis Enhancement

The two best Kemp eliminases from the library screening reported here share the presence of a bulky hydrophobic residue at the second position after the carboxyl terminus. It appears reasonable to assume then that this hydrophobic residue establishes interactions that stabilize the transition state of the reaction and enhance catalysis. In order to explore this possibility, we applied AlphaFold2 [39,40] to the prediction of the 3D-structure of our two best Kemp eliminases (Figure 5). We actually carried out the prediction for sequences in which the function-generating mutation (tryptophan to aspartate at position 229 in the β-lactamase sequence) was omitted. The reason for this is that W229 in β-lactamases is in roughly at the same position as the bound transition state in the variants with the W229D mutation that show Kemp elimination activity. Therefore, possible interactions that stabilize the bound transition state in the W229D variants may be suggested by the corresponding interactions with the tryptophan at position 229 in structures in which the W229D mutation is not included. Indeed, such interactions between the tryptophan and the hydrophobic residue at the second position are clearly suggested by the AlphaFold predictions (Figure 5).

**Figure 5.**
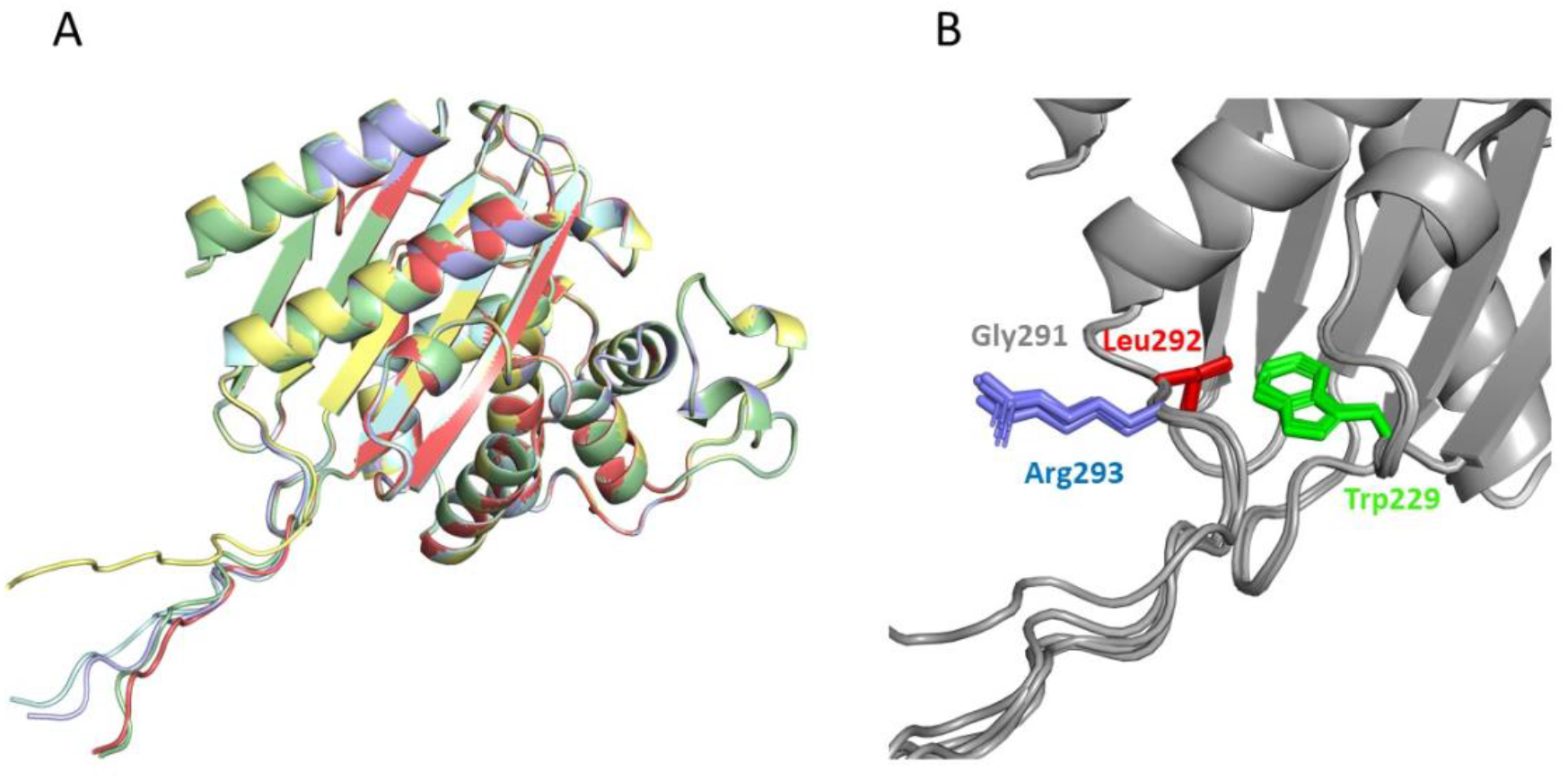
Prediction of the structure of the best Kemp eliminase obtained in this work by the program AlphaFold2 [39]. The prediction has been carried out with a sequence that has a tryptophan at position 229, *i.e*., the function-generating W229D mutation has been omitted. The reason for this is that the tryptophan at position 229 occupies a position close to that of the transition structure in the active Kemp eliminase (Figure 2A). Therefore, interactions with W229 in the predicted structure may conceivably correspond to interactions with the transition state that affect catalysis in the W229D variants. The top 5 predicted structures (**A**) are very close to the experimental structures shown in Figure 2, although the additional GLRG_3_SLEH_6_ polypeptide appears mostly extended and exposed to the solvent. Yet, a blow-up of the *de novo* active site region (**B**) reveals an interaction between the leucine of the polypeptide and the tryptophan at position 229 that could correspond to a kinetically relevant interaction with the transition state in the Kemp eliminase.

### 2.5 On the Possibility of Further Enhancements of Kemp Eliminase Activity

The catalysis enhancement levels achieved in this work are certainly substantial. Yet, it seems reasonable to ask whether they could be further enhanced using the same general approach. Of course, since we have only screened less than 10% of the different amino acid sequences spanned by the combinatorial library, it is obviously possible that better variants could be found by additional screening of the same library. More relevant is the fact that the library we have used is sharply focused to three positions. This allowed us to screen a substantial fraction of the library in a reasonable time. Still, it is conceivable that mutations at farther positions in the inserted polypeptide can also increase the rate of Kemp elimination. To explore this possibility, we prepared a new 8000-variant combinatorial library focused now at positions fourth, fifth and sixth after the carboxyl terminus (Figure 3C). Screening of about 250 variants from this library yielded results qualitatively similar to those obtained with the first library. Most of the variants displayed significant activity in the primary screening (Figure 3C) plausibly reflecting that the library samples the sequence space associated with a conformationally flexible polypeptide segment that likely remains largely exposed to the solvent in most cases. However, as was the case with the first library, a few of the variants were found to display substantially enhanced levels of catalysis for the Kemp elimination reaction (Figure 3C), a result that was confirmed by variant protein purification and determination of the Michaelis-Menten profiles and catalytic parameters (Figure 4A and Table 1). Certainly, the catalysis enhancement obtained on the basis of a limited screening of this second library is smaller than that afforded by the GLR variant from the first library. Yet, it is clear that combined screening of the positions included in the libraries is likely to lead to additional increases in *de novo* catalysis (work in progress).

## 3. Discussion

The generation of *de novo* enzymes (*i.e*., enzymes with completely new active sites) is one of the major unsolved problems in protein engineering [32]. Besides the obvious biotechnological implications, the results of *de novo* enzyme engineering studies may also have immediate implications for our understanding of the origin of life. Most of the chemical reactions of life are extremely slow in the absence of enzyme catalysis and several analyses support that diverse and specialized enzymes were already present in the last universal common ancestor [41,42]. It would seem reasonable to assume that efficient molecular mechanisms for the *de novo* emergence of enzymes and their subsequent optimization must exist. However, such efficient mechanisms are not apparent in the efforts of protein engineers to develop completely new enzyme functionalities. Only a limited number of *de novo* enzymes have been reported to date [32]and several of the most recent success stories in this field involve the recruitment of metals or metal-containing cofactors, which already provide by themselves some starting level of catalysis. However, only about 30% of enzymes are metalloenzymes [43,44]and mechanisms for the emergence of new enzymes that do not rely on metal recruitment remain poorly understood.

Furthermore, most engineered *de novo* enzymes display very low activities and many rounds of laboratory directed evolution, a highly time-consuming procedure, are often required to bring their catalysis to levels similar to those of modern natural enzymes [37,45-47].

Starting with the work of Tawfik, Baker and coworkers published in 2008 [48], Kemp elimination has been extensively used as a benchmark of *de novo* enzyme engineering. The reaction can occur through a base-catalysed mechanism, as shown in Figure 1A, and it is in fact considered as a model for proton abstraction from carbon, a fundamental process in chemistry and biochemistry. It can also occur through a redox mechanism and highly active *de novo* enzymes based on the recruitment of the heme cofactor for Kemp elimination catalysis have been recently reported [49,50]. Here, however, we are concerned with “traditional” Kemp elimination achieved through proton abstraction by a catalytic base (Figure 1A).

The best (at least in terms of k_cat_) base-promoted Kemp eliminase to date was reported by Hilvert and coworkers in 2013 [36] and displays a catalytic efficiency of k_cat_/K_M_=2.3·10^5^ M^−1^·s^−1^ and a turnover of k_cat_=700±60 s^−1^, values that compare favourably with those for the average modern natural enzyme (k_cat_/K_M_ about 10^5^ M^−1^·s^−1^ and turnover number k_cat_∼10 s^−1^; [35]). Remarkably, the catalytic parameters for the best Kemp eliminase found in this work, k_cat_/K_M_=(2.0±0.1)·10^5^ M^−1^·s^−1^ and k_cat_=635±59 s^−1^, agree with those for the best Kemp eliminase previously reported in the literature. These two efficient *de novo* Kemp eliminases are the result, however, of quite different design approaches and screening efforts. Hilvert’s Kemp eliminase was the outcome of 17 rounds of directed evolution from a rationally-designed *de novo* enzyme with a low, but significant level of Kemp eliminase activity [36]. The starting background for this extensive screening effort was the result of a complex iterative procedure in which a failed (*i.e*., inactive) initial computational design was rescued on the basis of amino acid replacements suggested by analyses of molecular dynamics simulations and 3D crystallographic structures [51]. By contrast, the best Kemp eliminase reported here started with a minimalist design targeting a conformationally flexible region in an ancestral β-lactamase scaffold [33]. A single mutation (W229D) thus generated a significant level of Kemp elimination activity which could be immediately enhanced by a second mutation (F290W) at the *de novo* active site. The Kemp elimination activity of this W229D/F2900W variant could be increased though computationally-focused, ultra-low-throughput screening [34].

Specifically, screening of only 20 active-site variants at the top of the stability ranking predicted by the FuncLib approach [52] led to a substantial activity improvement. Finally, in this work we have achieved further catalysis enhancement on the basis of the screening of only about 800 variants from a library that samples sequence space of a short polypeptide segment engineered at the active site region.

Our success in arriving at an efficient base-promoted Kemp eliminase on the basis of a rather modest screening effort has immediate implications for the engineering of *de novo* enzymes. First of all, the catalysis-enhancing mutations identified in this and our previous work [33,34] do not increase the complexity of the catalytic machinery, which remains a simple proton abstraction by a catalytic base, as established by the function-generating mutation W229D. Rather, they appear to optimize interactions that stabilize the transition state for the reaction ([34,37] and this work). Overall, it emerges that large enhancements in *de novo* enzyme catalysis can be achieved through the fine-tuning of intramolecular interactions in the active site region. Secondly, this work demonstrates that a short polypeptide segment inserted near the new active site has the capability of generating such catalysis-enhancing interactions. The approach has several obvious advantages. The inserted polypeptide segment will be in principle flexible and exposed to the solvent to a substantial extent and it will be thus unlikely to lead to disruptive interactions that compromise proper protein folding. Also, being a short segment, the associated sequence space can be efficiently sampled on the basis of a moderate screening effort.

Of course, the proof of principle provided in this work takes advantage of the fact that the carboxyl terminus in β-lactamases is close to the location of the new active site and, therefore, it can be easily used as the point of attachment of the new segment. This does not mean, however, that the extra-segment approach necessarily requires that the *de novo* active site is engineered near the carboxyl (or the amino) terminus of the protein. In fact, the new active site could be placed in any appropriate region of the protein, *i.e*., where design is successful, and then the carboxyl (or amino) terminus could be moved to a position close the new active site by engineering a protein variant with a suitable circular permutation [53].

## 4. Methods and Materials

### 4.1 Site-saturation Mutagenic Libraries

Library preparation was performed using the QuikChange Lightning PCR method (Agilent # 210518). Three positions were simultaneously saturated with the mutagenic primers described in Supplementary Table S2 in order to generate a combinatorial library comprising 8000 variants. The recombinant plasmid pET24-GNCA4-12-5G_HT containing the gene of the background variant was used as template. The amplification reaction contained 5 μL of 10× QuikChange Lightning Buffer, 1 μL of dNTP mix, 1.5 µl of QuikSolution reagent, 1.25 μL of primers (10 μM each mix), template plasmid (50 ng), 1 µl of QuikChange Lightning Enzyme and water to a final volume of 50 µL. The conditions for the PCR were as follows: 1 cycle at 95 °C for 2 min, 18 cycles of denaturation at 95 °C for 20 s, annealing at 60 °C for 10 sec, and extension at 68 °C for 3.5 min. The final extension step was carried out at 68 °C for 5 min. The PCR products were digested with DpnI. Two μL of the commercial DpnI solution were added to the PCR sample and the solution was incubated at 37 °C for 5 min. Afterwards, 2 µL of this solution were used to transform *E. coli* XL10-Gold Ultracompetent cells (45 μL) with a heat pulse. Subsequently, cells were suspended in 1 mL of SOC medium, incubated for 1 h at 37 °C, and plated on LB-agar containing 100 μg/mL kanamycin. To test the quality of the library, ten clones were randomly selected, their plasmids were extracted, and the gene of the β-lactamase variants was sequenced (Sanger).

### 4.2 Library Screening

*E. coli* BL21 (DE3) cells were transformed with the plasmid containing the mutant libraries or, as a control the variant used as background for library construction (Figure 3A), plated on LB-Kan agar and grown for 16 h at 37 °C. Individual colonies were picked and transferred into 44 mm deep well plates containing LB-Kan medium (0.2 mL) using a Pickolo colony picker with a Freedom EVO 200 robot from TECAN (Männedorf, Schweiz). Each plate contained an internal standard with the variant used as library background (column 7, rows A to H) and a negative control (column 1, row H). These master plates were incubated at 37 °C with shaking at 250 rpm. After 16 h, clones from this pre-culture were inoculated (using a cryo-replicator CR1000 from Enzyscreen, Haarlem, Netherlands) into deep well plates with fresh LB-Kan (lysate plates). After 4 hours of incubation at 37 °C, 250 rpm, LB-Kan with IPTG was added. The plates were incubated at 25 °C with shaking at 250 rpm for 16 h. The plates were centrifuged at 3000 g, the medium was discarded and the cell pellets were frozen at −80 °C. After ∼2 h the frozen cell pellets were re-suspended in HEPES 100 mM, pH 7. After 60 min at 25 °C the lysates were centrifuged at 3000 g and the supernatant was used for the Kemp eliminase assay.

### 4.3 Kemp Eliminase Assay for Library Screening

100 µL of the supernatant from lysate plates were transferred with the help of a liquid handler station (Freedom EVO 200, TECAN, Männedorf, Schweiz) to the reaction plates. The initial activities and residual activities values were determined by adding 100 μL of 100 HEPES 100 mM buffer pH 7.0 containing 0.25 mM 5-nitrobenzisoxazole. Plates were stirred briefly and the absorption at 380 nm (extinction coefficient of 15 800 M^−1^ cm^−1^) was followed in kinetic mode in the plate reader (Tecan M200 Infinite Pro Microplate Reader, Männedorf, Schweiz). The values were normalized against the average value corresponding to the background variant used for library construction. The best variant according to this determination were subsequently prepared and tested on pure form as described below.

### 4.4 Protein Expression and Purification

The various β-lactamase variants studied in this work were prepared and purified as described previously [33,34]. Briefly, genes cloned into a pET24-b vector with resistance to kanamycin were transformed into *E. coli* BL21(DE3) cells. The proteins were prepared with a His-tag and purified by affinity chromatography. Stock solutions for activity determinations and physicochemical characterization were prepared by exhaustive dialysis against the desired buffer.

### 4.5 Determination of Profiles of Rate versus Substrate Concentration for Kemp Eliminases

Rates of Kemp elimination were determined by following product formation by measuring the absorbance at 380 nm. An extinction coefficient of 15 800 M^−1^ cm^−1^ was used to calculate rates from the initial linear changes in absorbance with time. Most measurements were performed at 25ºC in HEPES 10 mM NaCl 100 mM pH 7 and 1% acetonitrile, although determinations in the same buffer at pH 8.5 were also performed for the best Kemp eliminase found in this work. The concentration of acetonitrile stated (1%) is the final concentration and it is the same for all the experiments. Variable amounts of acetonitrile were added to solutions with different substrate concentrations to ensure constancy of the final acetonitrile after adding different volumes of the substrate stock solution in acetonitrile. The absorbance increase at 380 nm was linear during the measurement and was recorded during a 15s interval, ensuring constant initial velocity conditions. All activity measurements were corrected by a blank performed under the same conditions. Still, the determined rates were always clearly above the blanks. Profiles of rate *versus* substrate concentration were used to calculate the values of the catalytic efficiency (k_cat_/K_M_), the turnover number (k_cat_) and the Michaelis constant (K_M_) by fitting the Michaelis-Menten equation to the experimental data, as we have described previously [33,34]. For each of the Kemp eliminases characterized in detail in this work, we actually obtained and analysed three different profiles of rate *versus* substrate concentration starting with at least two different protein preparations.

### 4.6 Protein Stability Determinations

Thermal stabilities of all the β-lactamase variants studied in this work were assessed by differential scanning calorimetry in HEPES 10 mM NaCl 100 mM pH 7 using a VP (Valerian Plotnikov) Capillary DSC (Microcal, Malvern) following protocols well established in our laboratory [7]. A typical calorimetric run involved several buffer– buffer baselines to ensure proper equilibration of the calorimeter followed by runs with several protein variants with intervening buffer–buffer baselines. A single calorimetric transition was observed in all cases. We have used the denaturation temperature, defined as the temperature corresponding to the maximum of the calorimetric transition, as a metric of protein stability.

### 4.7 Protein Structure Prediction

Protein structure prediction of variants obtained by directed evolution was performed using the ColabFold notebook implemented in Google Colab [40], based in the AlphaFold2 algorithm [39]. Sequences were used as inputs and structure prediction was performed with default parameters. PyMOL (PyMOL Molecular Graphics System, Version 2.4.1 Schrödinger, LLC) was used to visualize the predicted models and inspect the role of the different mutations.

## Supplementary Materials

The following supporting information can be downloaded at www.mdpi.com/xxx/s1, Figure S1: Michaelis-Menten profiles for engineered Kemp eliminases. Table S1: Catalytic parameters for Kemp eliminases. Table S2: Primers used for saturation mutagenesis.

## Author contributions

L.I.G.-R. and V.A.R. performed the library screening experiments, purified the selected β-lactamase variants and determined their biophysical features. M.A. provided essential input regarding library design and the interpretation of library screening results. L.I.G.-R. carried out the structural analyses of Kemp eliminases using AlphaFold2. V.A.R. and J.M.S.-R. designed the research. J.M.S.-R. wrote the first draft of the manuscript. All authors discussed the manuscript, suggested modifications and improvements and contributed to the final version.

## Data availability statement

All relevant data are included in the manuscript and its Supplementary Materials file.

