## Supplemental Figures and Tables for "Efficient base-catalysed Kemp elimination in an engineered ancestral enzyme"

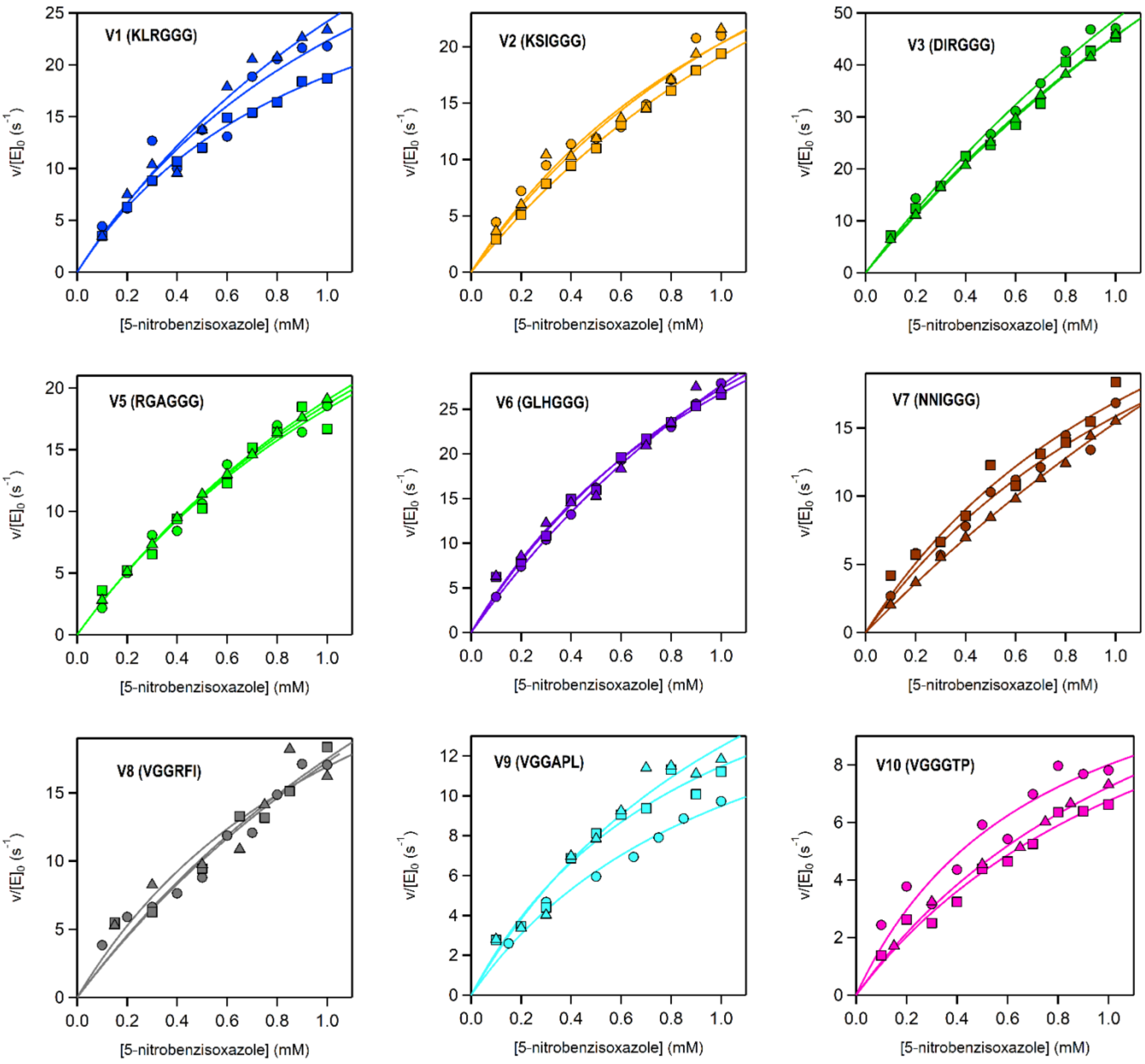


**Figure S1**. Michaelis-Menten profiles of the three independent determinations for the top variants from the primary library screening at pH 7. The sequences at the relevant section of the included polypeptide are shown for each variant.

**Table S1**. Catalytic parameters for the cleavage of 5-nitrobenzisoxazole at pH 7 (HEPES 10 mM NaCl 100 mM) and 1% acetonitrile and 25 ºC catalyzed by the engineered and evolved versions of Precambrian β-lactamases. Values of catalytic parameters derived from the fitting of the Michaelis-Menten equation are given for each of the three independent replicates. Errors are represented as the standard deviation derived from the fitting. The sequences of each variant at the randomly mutagenized regions in the libraries are displayed shown.

| Variant | Sequence[a] | $k_{cat}$ ($s^{-1}$) | $K_M$ (mM) | $k_{cat}/K_M$ ($s^{-1}$ $M^{-1}$) |
|---|---|---|---|---|
| V4 (pH 8.5) | GLRGGG | 703 ± 104 | 6.67 ± 0.65 | $(1.9 \pm 0.04) \times 10^5$ |
| | | 606.4 ± 181 | 3.05 ± 1.14 | $(2.0 \pm 0.1) \times 10^5$ |
| | | 596.9 ± 164 | 2.71 ± 0.85 | $(2.2 \pm 0.2) \times 10^5$ |
| V4 (pH 7.0) | GLRGGG | 299.0 ± 99.8 | 2.33 ± 1.02 | $(1.3 \pm 0.1) \times 10^5$ |
| | | 460.9 ± 53.3 | 4.54 ± 0.61 | $(1.02 \pm 0.02) \times 10^5$ |
| | | 462.6 ± 143.0 | 3.19 ± 1.22 | $(1.5 \pm 0.1) \times 10^5$ |
| V3 | DIRGGG | 206.8 ± 55.3 | 3.24 ± 1.07 | $(6.4 \pm 0.4) \times 10^4$ |
| | | 190.4 ± 54.4 | 3.18 ± 1.12 | $(6.0 \pm 0.4) \times 10^4$ |
| | | 211.4 ± 17.7 | 3.65 ± 0.37 | $(5.8 \pm 0.1) \times 10^4$ |
| V6 | GLHGGG | 93.3 ± 6.5 | 2.38 ± 0.22 | $(3.9 \pm 0.1) \times 10^4$ |
| | | 61.8 ± 7.4 | 1.31 ± 0.24 | $(4.7 \pm 0.3) \times 10^4$ |
| | | 69.1 ± 14.9 | 1.53 ± 0.48 | $(4.5 \pm 0.4) \times 10^4$ |
| V1 | KLRGGG | 53.8 ± 18.2 | 1.41 ± 0.72 | $(3.8 \pm 0.7) \times 10^4$ |
| | | 38.0 ± 2.6 | 1.01 ± 0.12 | $(3.8 \pm 0.2) \times 10^4$ |
| | | 70.4 ± 20.4 | 1.91 ± 0.76 | $(3.7 \pm 0.4) \times 10^4$ |

| | | | | |
|---|---|---|---|---|
| V2 | KSIGGG | 49.1 ± 12.8 | 1.42 ± 0.55 | $(3.5 \pm 0.5) \times 10^4$ |
| | | 60.9 ± 5.2 | 2.18 ± 0.25 | $(2.8 \pm 0.1) \times 10^4$ |
| | | 54.1 ± 14.3 | 1.66 ± 0.63 | $(3.3 \pm 0.4) \times 10^4$ |
| V5 | RGAGGG | 55.0 ± 12.5 | 1.95 ± 0.61 | $(2.8 \pm 0.2) \times 10^4$ |
| | | 52.3 ± 15.2 | 1.86 ± 0.75 | $(2.8 \pm 0.3) \times 10^4$ |
| | | 59.0 ± 1.9 | 2.10 ± 0.09 | $(2.81 \pm 0.03) \times 10^4$ |
| V8 | VGGRFI | 61.8 ± 27.0 | 2.58 ± 1.45 | $(2.4 \pm 0.3) \times 10^4$ |
| | | 60.8 ± 34.0 | 2.47 ± 1.81 | $(2.4 \pm 0.4) \times 10^4$ |
| | | 38.0 ± 18.2 | 1.24 ± 0.95 | $(3.1 \pm 0.9) \times 10^4$ |
| V7 | NNIGGG | 41.0 ± 10.6 | 1.59 ± 0.60 | $(2.6 \pm 0.3) \times 10^4$ |
| | | 40.4 ± 11.2 | 1.39 ± 0.58 | $(2.9 \pm 0.4) \times 10^4$ |
| | | 84.8 ± 14.1 | 4.51 ± 0.88 | $(1.9 \pm 0.1) \times 10^4$ |
| V9 | VGGAPL | 20.0 ± 3.0 | 1.11 ± 0.27 | $(1.8 \pm 0.2) \times 10^4$ |
| | | 22.2 ± 3.9 | 0.94 ± 0.28 | $(2.4 \pm 0.3) \times 10^4$ |
| | | 28.9 ± 7.6 | 1.31 ± 0.53 | $(2.2 \pm 0.3) \times 10^4$ |
| V10 | VGGGTP | 17.6 ± 2.4 | 1.43 ± 0.30 | $(1.2 \pm 0.1) \times 10^4$ |
| | | 13.9 ± 2.8 | 0.74 ± 0.28 | $(1.9 \pm 0.3) \times 10^4$ |
| | | 16.1 ± 3.5 | 1.39 ± 0.45 | $(1.2 \pm 0.1) \times 10^4$ |
| BACKGROUND | VGGGGG | 21.9 ± 4.0 | 2.35 ± 0.56 | $(9.4 \pm 0.6) \times 10^3$ |
| | | 11.3 ± 3.8 | 1.21 ± 0.65 | $(9.3 \pm 1.9) \times 10^3$ |
| | | 15.7 ± 2.3 | 1.56 ± 0.33 | $(1.0 \pm 0.1) \times 10^4$ |

**Table S2**. Sequences for the mutagenic primers used to saturate three simultaneous positions of the included polypeptide for the generation of the combinatorial libraries.

| COMBINATORIAL LIBRARY | PRIMER NAME | PRIMER SEQUENCE |
|---|---|---|
| Library 1 (VGGGGG) | VHH_F | gtggtggtggtggtggtgctcgagtgaNNNNNNNNN accacccacccacgccgccacaaccagacgc |
| | VHH_R | gcgtctggttgtggcggcgtgggtgggtggtNNNNNNNNN tcactcgagcaccaccaccaccaccac |
| Library 2 (VGGGGG) | HHH_F | gtggtggtggtggtggtgctcgagtgagccaccacc NNNNNNNNNccacgccgccacaaccagacgc |
| | HHH_R | gcgtctggttgtggcggcgtggNNNNNNNNN ggtggtggctcactcgagcaccaccaccaccaccac |
